# Chemoenzymatic labeling of DNA methylation patterns for single-molecule epigenetic mapping

**DOI:** 10.1101/2021.02.24.432628

**Authors:** Tslil Gabrieli, Yael Michaeli, Sigal Avraham, Dmitry Torchinsky, Matyas Juhasz, Ceyda Coruh, Nissim Arbib, Zhaohui Sunny Zhou, Julie A. Law, Elmar Weinhold, Yuval Ebenstein

**Author notes:** Correspondence: Yuval Ebenstein.; Elmar Weinhold.

## Abstract

DNA methylation, specifically, methylation of cytosine (C) nucleotides at the 5-carbon position (5-mC), is the most studied and among the most significant epigenetic modifications. Here we developed a chemoenzymatic procedure to fluorescently label non-methylated cytosines in the CpG context allowing epigenetic profiling of single DNA molecules spanning hundreds of thousands of base pairs. For this method, a CpG methyltransferase was used to transfer an azide to cytosines from a synthetic *S*-adenosyl-l-methionine cofactor analog. A fluorophore was then clicked onto the DNA, reporting on the amount and position of non-methylated CpGs. We found that labeling efficiency was increased two-fold by the addition of a nucleosidase that degrades the inactive by-product of the azide-cofactor after labeling, and prevents its inhibitory effect. We first used the method to determine the decline in global DNA methylation in chronic lymphocytic leukemia patients and then performed whole genome methylation mapping of the model plant *Arabidopsis thaliana.* Our genome maps show high concordance with published methylation maps produced by bisulfite sequencing. Although mapping resolution is limited by optical detection to 500-1000 base pairs, the labeled DNA molecules produced by this approach are hundreds of thousands of base pairs long, allowing access to long repetitive and structurally variable genomic regions.

## INTRODUCTION

DNA methylation is an epigenetic mark that plays a major regulatory role in transcription, gene regulation, and disease. 5-methylcytocine (5-mC) is conserved among plants and mammals, and its precise genomic patterns are crucial for development. In mammals, 5-mC occurs most often in the context of CpG dinucleotides and, in the human genome, most CpGs are highly methylated (between 70-80%). However, in specific locations, mainly high density CpG islands (CGI), they remain mostly non-methylated. The majority of CGIs (~70%) are located at gene promoters and the methylation status of these regions are known to regulate gene expression^1,2,3^. In cancer, large-scale changes in methylation levels are observed and can include both genome-wide hypomethylation as well as more localized hypermethylation, mainly of tumor suppressor genes that are often silenced in cancer^4,5,6,7,8^.

Methylation is established by a diverse family of methyltransferases (MTases). These enzymes utilize *S*-adenosyl-L-methionine (AdoMet) as the methyl donor, forming methylated DNA and *S*-adenosyl-L-homocysteine (AdoHcy) as a by-product^9^. Various approaches have been developed to specifically profile cytosine DNA methylation, both on a global level and at base pair resolution. The gold standard technique is bisulfite sequencing (called BS-seq (Cokus et al.^10^) or methylC-seq (Lister et al. ^11^)) depending on the methods used), which relies on chemical conversion of unmodified cytosines to uracil, while leaving methylated cytosines unconverted^10,12^. After PCR amplification and sequencing, the originally methylated cytosines are read as C while the unmodified cytosines are read as T. However, this method can suffer from several limitations, such as high DNA degradation and potential biases due to the amplification process. Short read sequencing also adds several drawbacks and mainly the limitation in characterizing large variable and repetitive regions, as well as population averaging of the data which masks cell-to-cell variation^13^. Single-cell bisulfite sequencing may profile individual methylomes^14–15^ but is still limited by short read sequencing and thus is not able to fully characterize highly repetitive regions. Additionally, the technique suffers from low sampling of the individual genomes, thus requiring sequencing the genomes of thousands of individual cells^16^.

Third generation sequencing approaches, including single-molecule, real-time (SMRT) sequencing (Pacific Biosciences Inc.) and nanopore sequencing (Oxford Nanopore Technologies Ltd.), are able to detect methylated cytosines directly^17–21^. However, the accuracy of epigenetic analysis on these platforms is still lacking and whole genome analysis is challenging^22^.

Whole-genome epigenetic profiling by optical genome mapping (Bionano Genomics Inc.) is a new addition to the epigenetic mapping toolbox. This technology is based on stretching long genomic DNA fragments for optical imaging. The DNA is labeled with two colors, one is used for aligning the molecules to the reference genome and the other reports on the epigenetic content of the molecules^23–25^. This technique outputs extremely long, single-molecule data (N50~200 kbp) that allows profiling of large variable and repetitive regions^26^.

Our group developed Reduced representation Optical Methylation mapping (ROM), as a method for the labeling and detection of non-methylated CpG sites using the bacterial MTase, M.TaqI, and a cofactor analogue^24^. Recently, this long-read methylation data was analyzed at a single-cell level for the first time. The methylation status of promoters and their distal enhancers, simultaneously imaged on the same long DNA molecules, served to accurately deconvolve cell-type mixtures and subpopulations within a sample^27^. The non-methylation labeling scheme relies on the transfer of a fluorescent molecule from a synthetically modified form of the native methylation cofactor AdoMet. The enzyme is "tricked" into transferring the extended chemical group with a fluorophore instead of the natural methyl group, resulting in fluorescence at non-methylated CpG sites. However, M.TaqI (recognition site: TCGA) only samples about 6% of human CpG sites and thus can fail to capture vital epigenetic information.

Here we present a chemoenzymatic labeling approach for the detection of non-methylated CpGs as a means for a global quantification of methylation and for genomic methylation mapping. This is achieved by utilizing the CpG-specific MTase M.SssI from the bacteria *Spiroplasma sp.* strain MQ-1 that naturally transfers a methyl group to the fifth position of cytosines in CpG dinucleotides^28,29^. While M.SssI has the potential to label all non-methylated CpGs, it is not able to utilize modified AdoMet cofactors. However, an engineered variant of M.SssI (Q142A/N370A) is able to transfer an azide group to non-methylated CpG sites by processing an azide-modified cofactor AdoYnAzide (Ado-6-azide). This engineered M.SssI (eM.SssI) was used by Kriukiene *et al.* for non-methylation sequencing^30^. However, the affinity capture on streptavidin-modified beads showed a 20-30% capture efficiency, indicating poor labeling yield which is insufficient for single-molecule optical mapping. To overcome the low labeling efficiency, we employed the enzyme 5-methylthioadenosine/*S*-adenosylhomocysteine nucleosidase (MTAN, EC 3.2.2.9) that catalyzes hydrolysis of the glycosidic bond in AdoHcy. Thus, MTAN degrades the inactive cofactor by-product, AdoHcy, formed from the natural cofactor AdoMet or synthetic cofactor analogues, like AdoYnAzide (Figure 1A). In general, AdoHcy is a well-known product inhibitor for MTases^31^. Lowering its concentration by addition of MTAN effectively drives the reaction towards increased labeling efficiency (Figure 1A). The azide-modified DNA can then be covalently labeled by a dibenzocyclooctyne-cy5 (DBCO-cy5) fluorophore in a strain-promoted azide-alkyne cycloaddition (SPAAC, copper-free click chemistry reaction), to specifically label non-methylated CpG sites. Thus, the increased efficiency of the eM.SssI / MTAN combination enabled generation of optical maps that present genome-wide epigenetic status of CpG sites with single-molecule sensitivity.

**Figure 1.**
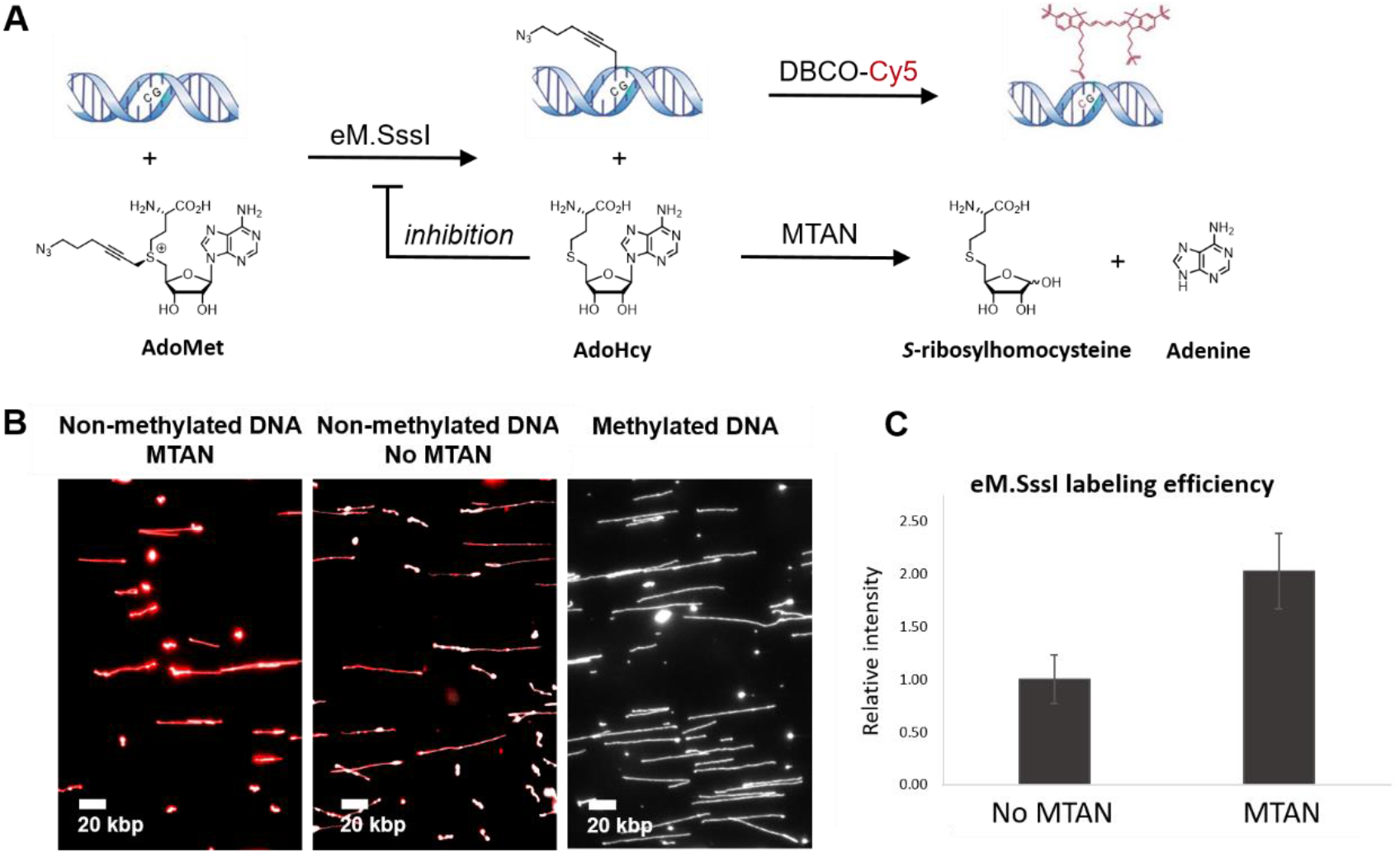
**A**. eM.SssI catalyzes the addition of AdoYnAzide to non-methylated CpGs. DBCO-cy5 is then attached by click chemistry to the azide-modified CpGs. Following the transfer of the azide-containing side chain by eM.SssI, the remaining cofactor by-product AdoHcy serves as a substrate for MTAN that hydrolyzes it to adenine and *S*-ribosylhomocysteine. In the absence of MTAN, AdoHcy accumulates and can bind to eM.SssI, inhibiting its activity. **B**. A representative field of view of non-methylated λ DNA labeled with eM.SssI, in the presence (Left) or absence (Middle) of MTAN. Methylated DNA was labeled as control (Right). Gray-DNA backbone, red-non-methylated CpG labels. **C**. Global quantification (calculated by dividing the total intensity of the CpG label by the total length of the DNA) of non-methylated CpGs labelled with eM.SssI, with or without MTAN.

Here, we apply this labeling strategy to human peripheral blood mononuclear cells (PBMCs), emphasizing the ability to quantify the methylation status of CpG sites in healthy vs. cancer patients with high sensitivity. We further demonstrate whole-genome methylation mapping of the model plant *A. thaliana*. Our approach allows simultaneous genetic and epigenetic characterization of long streches of DNA at the single molecule level, and potentially permits studying variations between single cells.

## RESULTS AND DISCUSSION

### Efficient labeling of non-methylated CpGs

Single-molecule epigenomic mapping requires highly efficient labeling in order to extract the maximum information from the studied sample. We hypothesized that during labeling by eM.SssI the cofactor by-product AdoHcy could reduce the overall labeling efficiency by occupying the cofactor binding pocket and preventing additional labeling cycles (Figure 1A). Thus, we sought to enzymatically degrade this by-product by the addition of recombinant MTAN^32^. To test the effect of MTAN on the labeling efficiency of eM.SssI, we labeled non-methylated λ DNA using eM.SssI, AdoYnAzide, and DBCO-cy5, with or without MTAN (Figure 1B & Figure S1). Figure 1C shows the relative fluorescence intensity of λ DNA labeled with or without MTAN. Average labeling intensity increased two-fold with the addition of MTAN to the reaction (Figure 1C, Figure S2) and reached a level that enables reliable single-molecule analysis and epigenetic genome mapping.

### Global methylation quantification

One of the hallmarks of cancer is global reduction in genomic methylation levels^33^. As a first step in evaluating our optical mapping technique for epigenetic profiling, we quantified the genome-wide methylation level of PBMCs from a patient diagnosed with chronic lymphocytic leukemia (CLL) and a healthy donor. Both healthy and patient DNA were stretched in nanochannel array chips, imaged, and the fluorescence from the non-methylated CpG labels was detected (Figure 2A). Fluorescence intensity along the labeled DNA molecules was automatically measured to determine the relative difference in labeling^34^. The relative global methylation level was measured by summing the overall red intensity (*i.e.* non-methylated cytosines in the CpG context) along the DNA molecules, normalized to the total DNA length measured for each sample (Figure 2B). When comparing genomic DNA from a patient and a healthy donor, the cancer cells show a global reduction in methylation levels indicated by a higher fluorescence signal along the DNA (Figure 2B). Specifically, the signal intensity from non-methylated CpGs increased by 2.3-fold in the CLL sample compared to the healthy control, indicating global hypomethylation in the CLL sample as previously reported (supplementary files 2&3)^35^. Incorporation of our novel chemoenzymatic labeling method into single molecule optical mapping allows us to distinguish global DNA methylation differences between samples, opening up potential application of this technique for DNA methylation research and medical diagnostics.

**Figure 2.**
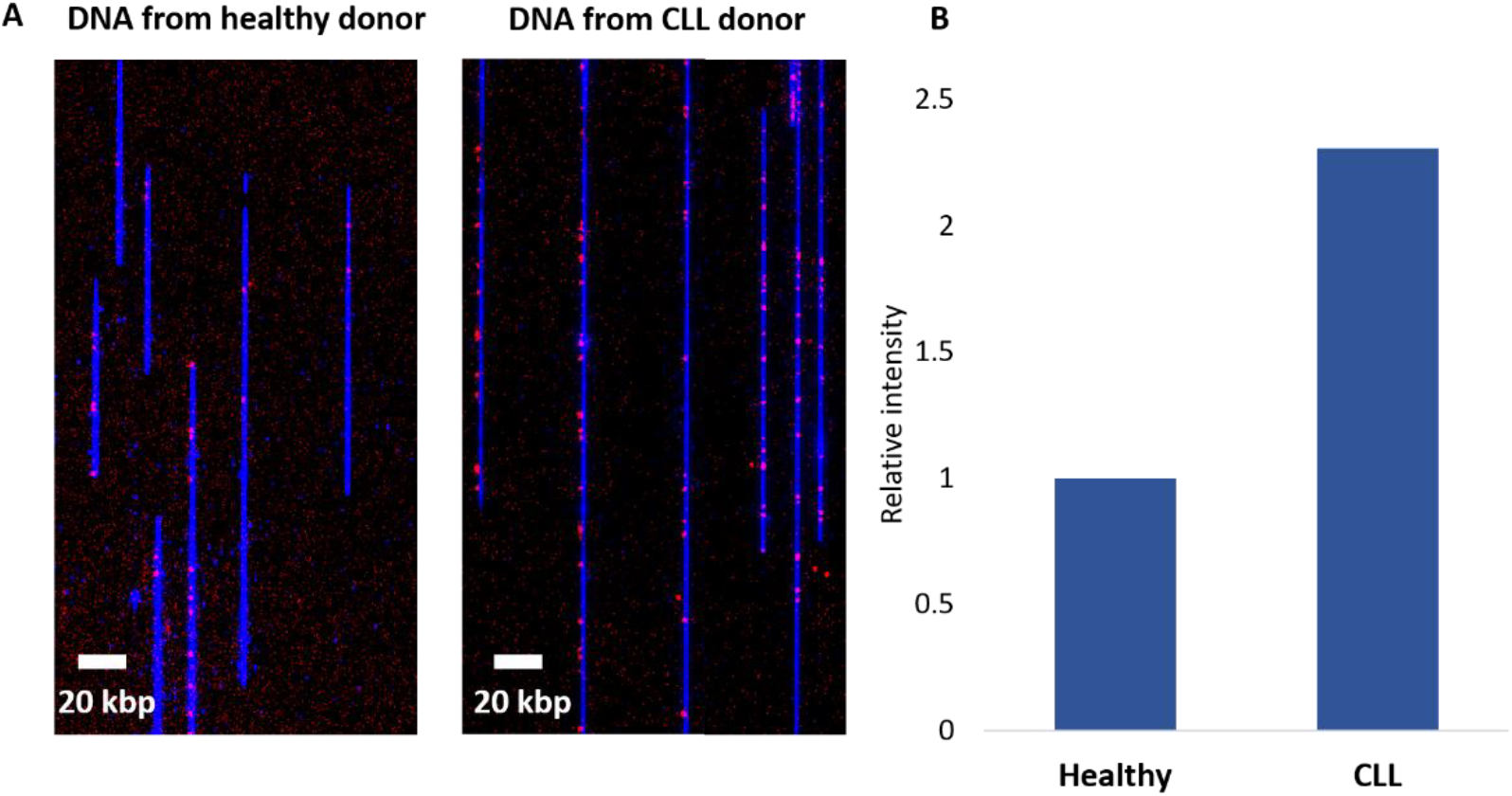
Methylation quantification in human peripheral blood mononuclear cells (PBMCs) from a healthy and CLL patient donor. **A**. Representative images of stretched DNA in nanochannel array chips from both samples: DNA backbone in blue and epigenetic labels in red (more red signal denotes less DNA methylation at CpG sites). **B**. Global quantification of the methylation signal intensity along the DNA molecules. Intensity of the healthy sample was normalized to 1.

### Genomic mapping of DNA methylation

We next attempted to map the CpG methylation profile in the well-characterized genome of the model plant, *Arabidopsis thaliana*. This genome is composed of five chromosomes with a total size of ~135 Mbp. The relatively small genome allowed us to test mapping feasibility while avoiding the cost and computational complexity of analyzing the human genome. In *A. thaliana* DNA methylation commonly occurs in all sequence contexts (CG, CHG and CHH, where H=A, T or C)^10,12^ and the presence of methylation at transposons and repeats, as well as some protein coding genes, is associated with gene silencing and genome stability^12,36,37^. *A. thaliana* DNA was extracted, non-methylated CpGs were labeled using our modified M.SssI system, and genomic mapping was performed by using nickase labeling to create a barcode of specific sequence motifs. The dual-labeled DNA molecules were then electro-kinetically forced into nanochannels and imaged on the Bionano Genomics (Inc) Irys system, allowing the detection of fluorescent labeling patterns (Figure 3A and 3B). We detected 34,866 DNA molecules longer than 100 kbp and 28,917 were successfully aligned to the TAIR10 *A. thaliana* reference genome (Figure S3). The analysis resulted in an average mapping depth of 47x, covering ~97% of the genome with a minimum coverage of 15 molecules. The intensity profile of the epigenetic labels for each molecule was created by Irys Extract^38^, followed by calculation of normalized average intensity across all molecules aligned to the same region (Figure 3C).

**Figure 3.**
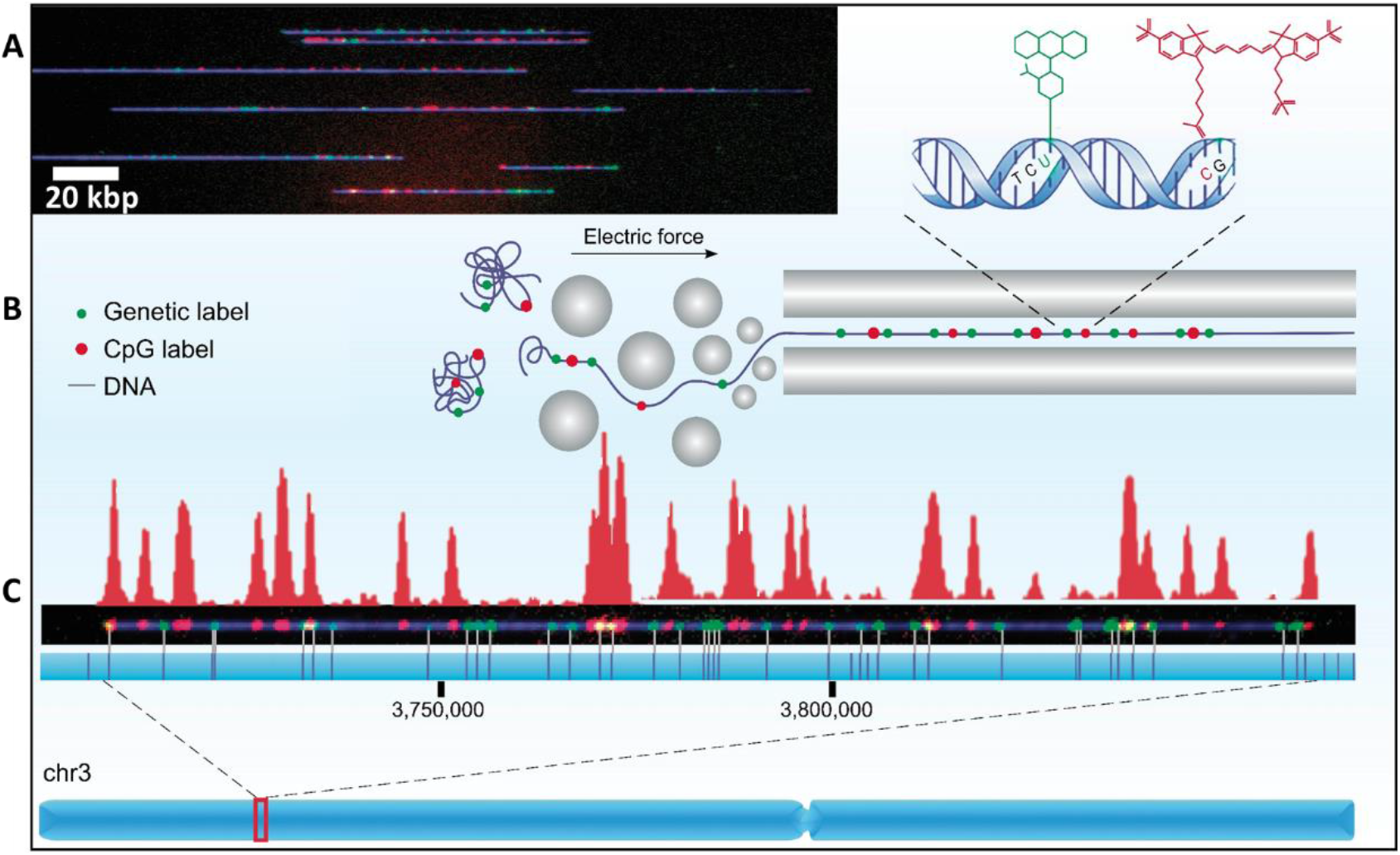
Optical methylation mapping scheme. **A**. A representative image of stretched DNA molecules in a nanochannel array. DNA backbone in blue (YOYO-1), genetic labels in green (Atto-532) and non-methylated CpG labels in red (cy5). **B**. Schematic representation of the fluorescently labeled molecules unraveled and extended in a nanochannel array by using an electric field. **C.** A 150 kbp DNA molecule aligned to chromosome 3 by the green genetic labels. The vertical lines on the blue strip represent the theoretical positions of genetic labels in the reference genome. The fluorescence intensity pattern representing the levels of non-methylated CpG sites along the molecule is presented in red above the molecule.

### Optical methylation mapping is well correlated with bisulfite sequencing

To further evaluate methylation mapping, a genome-wide correlation of publicly-available bisulfite sequencing data^36^ with epigenetic optical mapping was calculated. Since our method captures signal from non-methylated CpGs, we present the genome-wide bisulfite sequencing data as the non-methylated fraction of CpGs. To overcome scaling and resolution differences between the two methods, each non methylated CpG from the bisulfite sequencing data was padded +/−500 bp and summed across the genome in 10 kbp bins to generate a “methylation score” that corresponds to the amount of non-methylated CpG sites in each bin (*i.e.* a higher score equates to more non-methylated CpG sites,). This binned data was then compared to the average intensity profile of optical epigenome mapping. First, optical data was normalized for coverage variations by dividing the summed local intensity by the local coverage to obtain a fluorescence intensity score in 1 kbp bins across the genome. The normalized intensity score in each bin was then divided by the global maximum intensity (e.g. the 1 kbp bin with the highest fluorescence intensity score), setting the relative intensity data between 0-1. As presented in Figure 4A, the optical mapping data nicely correlate with the genome-wide bisulfite sequencing data with a Spearman correlation coefficient of 0.745 (supplementary file 4). In addition, a global view of the *A. thaliana* genome showing the levels of non-methylated CpG sites illustrates the correlation between the optical mapping and genome-wide bisulfite sequencing data (Figure 4B). The centromeres of the *A. thaliana* genome are enriched in methylated CpGs and are thus seen as dips in the optical graphs at the centromeric positions (low signal denotes high methylation, Figure 4B). The lower panel of Figure 4B shows a zoomed-in region in Chr1 illustrating the positive correlation between the two methods. The finer details in the blue track highlights the higher resolution of the genome-wide bisulfite sequencing data. Nevertheless, the long molecules observed by optical mapping (>100 kbp) allowed us to assemble a reliable and highly contiguous epigenetic profile of the *A. thaliana* genome.

**Figure 4.**
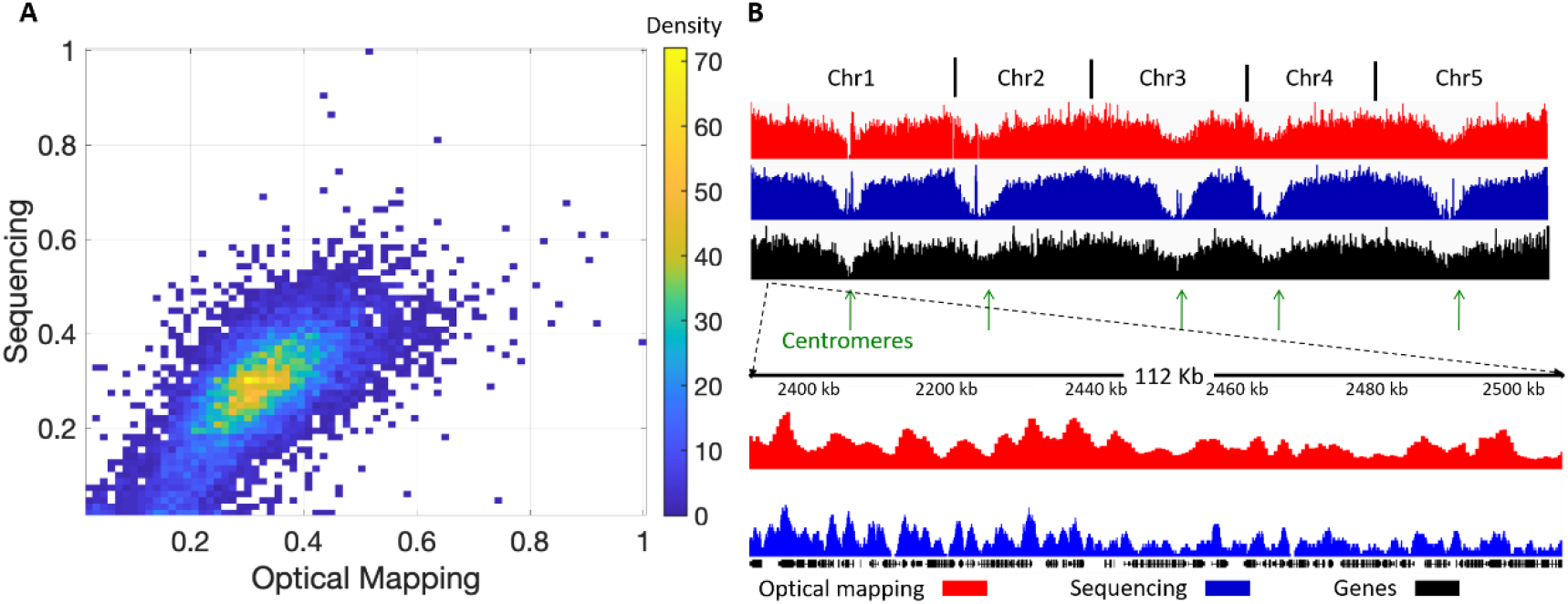
Global correlation between genome-wide bisulfite sequencing and methylation optical mapping. **A**. Density scatter plot representing the global correlation between genome-wide bisulfite sequencing and optical mapping in 10 kbp bins. **B**. Global view of the optical methylation profiles (relative intensity) in red and the genome-wide bisulfite sequencing data (methylation score) in blue, both showing the levels of non-methylated CpG sites. For reference, the genes are shown in black and the centromeres are indicated by green arrows. The bottom panel presents a zoomed-in view of a selected region.

### Optical patterns correlate with genome-wide bisulfite sequencing at siRNA clusters and gene bodies

In *A. thaliana*, DNA methylation at transposons and repeats is guided by short interfering RNAs (siRNAs) that recruit the DNA cytosine MTase, DOMAINS REARRANGED METHYLTRANSFERASE 2 (DRM2), *via* protein/RNA complexes^39,40^. Since the 24nt siRNAs guide where DNA methylation occurs in the plant genome, including methylation in the CpG context, we checked if the CpG methylation patterns from bisulfite sequencing data correlates as expected with the optical data at sites that produce 24nt siRNAs (*i.e.* sites producing 24nt siRNAs should have lower levels of non-methylated CpG sites compared to surrounding regions). Methylation scores from genome-wide bisulfite sequencing data (supplementary file 5) and relative intensity values from optical mapping data were calculated 10 kbp upstream and downstream relative to the 24nt siRNA clusters (position 0)^41^. These data were combined into a single plot representing the methylation profile around these regions (Figure 5A). The methylation score from bisulfite sequencing data (blue) shows a sharp dip around position 0, indicating highly methylated DNA around 24nt siRNA clusters. The relative intensity values from the optical mapping data (red) correlates well with this trend, although at lower resolution, as expected from the optical diffraction limit.

**Figure 5.**
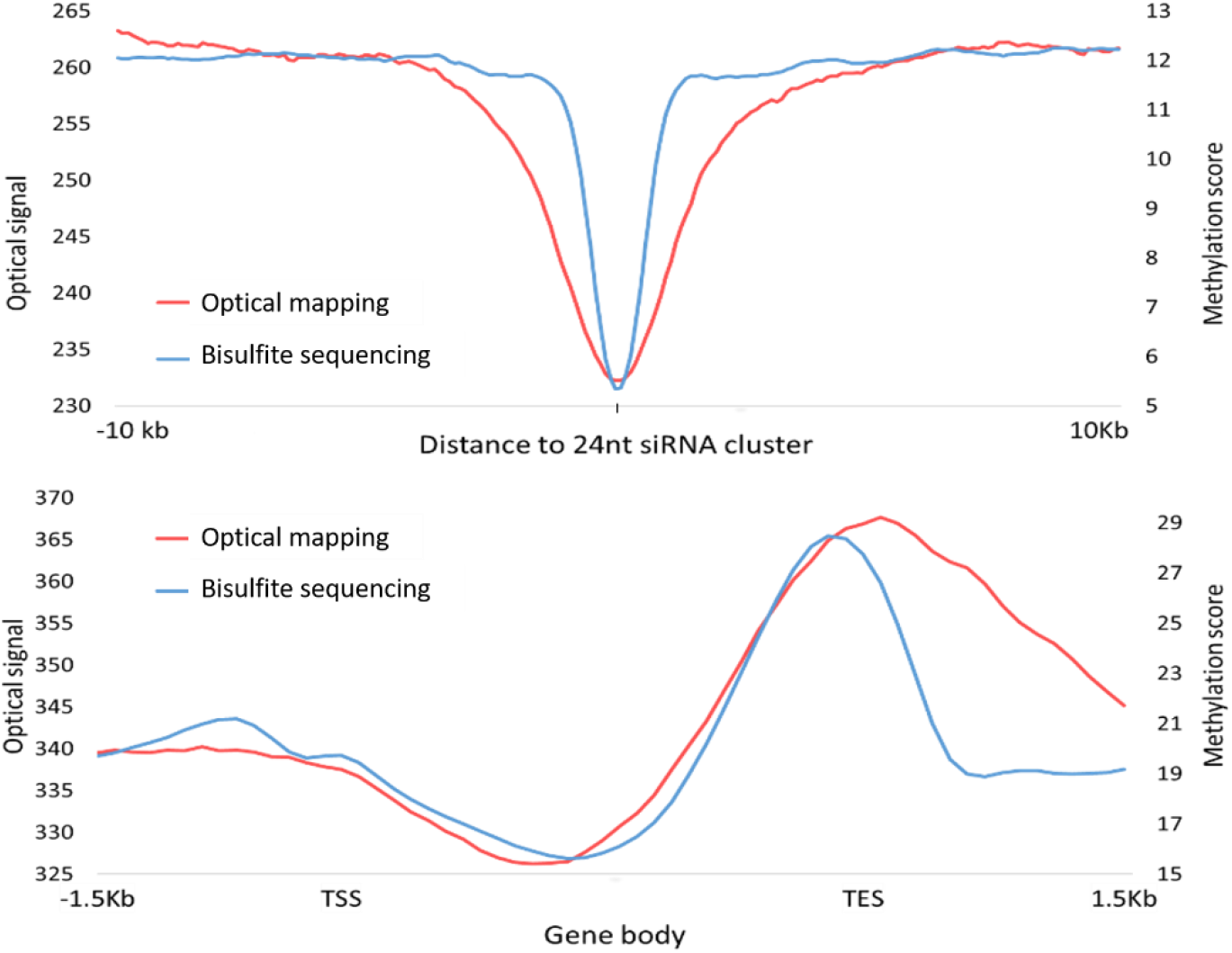
Methylation profiles at 24nt siRNA producing loci and across gene bodies. Genome-wide bisulfite sequencing data in blue & optical mapping data in red. **A**. 24nt siRNA loci. **B**. Methylation profile across gene bodies and 1.5 kbp upstream and downstream to the transcription start site (TSS) and the transcription end site (TES), respectively.

As negative controls, we examined optical mapping data over primary miRNA loci and several trans-acting siRNA loci (TAS loci) which are not related to DNA methylation (Figure S4, supplementary file 5). As expected, the intensity profile across the miRNA precursor and TAS loci did not have a specific pattern around the cluster center.

In addition to DNA methylation in all sequence contexts at transposons and repeats, some gene bodies are methylated specifically in CpG context in the *A. thaliana* genome and this methylation is distributed away from the 5’ region of genes^10^. Thus, we also compared the methylation profiles across gene bodies from genome-wide bisulfite sequencing and optical mapping (supplementary file 6). Figure 5B illustrates the profiles across the gene bodies and 1.5 kbp upstream of the transcription start site (TSS) and downstream from the transcription end site (TES). Again, the two datasets show the expected modulation in methylation along the gene, with sequencing data displaying higher resolution (as seen downstream of the TES). While lacking in resolution, optical mapping has the advantage of visualizing intact genes as well as extremely long DNA molecules that are able to capture both large-scale methylation patterns, and methylation across highly variable regions that are difficult to map by short-read sequencing.

## CONCLUSIONS

In this study we use the DNA methyltransferase eM.SssI and a synthetic cofactor together with the nucleosidase MTAN to fluorescently label non-methylated CpG dinucleotides. The addition of MTAN to the reaction dramatically increased the labeling efficiency of eM.SssI, making the method compatible with single-molecule analysis. Application of this labeling approach to optical genome mapping enables genome-wide methylome profiling, offering long-range information that can span long variable regions and repeats^23,24^. We note that this concept may be similarly utilized to map other genomic features such as other epigenetic modifications and DNA damage lesions ^23,24,42–45^.

Global methylation changes can be quantified by integrating the optical methylation signal for a given amount of DNA (Figure 2). Assessment of whole genome methylation levels may serve as a diagnostic tool that can distinguish between patients with healthy or malignant cells. It is long known that patients with B-cell chronic lymphocytic leukemia (CLL) experience hypermethylation in specific genes^46^ while their global methylation levels decrease^35^. This hypomethylation can be detected by the presented optical labeling scheme with very high sensitivity, emphasized by the distinct differences in labeling between the healthy and CLL patient (Figure 2B).

The generation of whole genome methylation maps by means of optical mapping was demonstrated here on the model plant *A. thaliana* and found to correlate well with genome-wide bisulfite sequencing data. As many plant genomes are highly repetitive, optical mapping is becoming a routine method for genomic characterization in plants^47–49^. By utilizing this new approach, it is now possible to access methylation information in repetitive or otherwise challenging regions. While optical mapping resolution is limited by the optical diffraction limit to around 1 kbp, we increased the effective resolving power of the technique by using fluorescence intensity in addition to genomic location. Thus, CpG-rich regions, where more than one CpG site resides within the optical spot, can now be characterized by signal intensity that reports on labeling density, *i.e.*, the actual number of labeled CpG sites at a given fluorescent spot. Future extension of the method to the human genome may facilitate early diagnosis of malignant transformations related to DNA methylation.

## METHODS

### Human Subjects

The healthy donor sample used in this study was collected with informed consent for research use and approved by the Tel-Aviv University and Meir medical center ethical Review Boards, in accordance with the declaration of Helsinki. The PBMCs from a CLL donor were purchased from BioServe.

### High-molecular-weight DNA extraction

High molecular weight (HMW) DNA was extracted for optical mapping from 1 g of 6-8 weeks old whole plant tissue, before flowering using the BioNano Genomics Fix’n’Chop protocol with some modifications. Briefly, no formaldehyde was used, chopping was done only with lab blender and no razor blade was used, 7.5% of 2-Mercaptoethanol was used. Following the final wash, nuclei pellet was resuspended in cell suspension buffer (CHEF mammalian DNA extraction kit, Bio-Rad) and incubated at 43 °C for 10 min. 2% low melting agarose (CleanCut agarose, Bio-Rad) was melted at 70 °C followed by incubation at 43°C for 10 min. Melted agarose was added to the resuspended cells at a final concentration of 0.75% and mixed gently. The mixture was immediately cast to a plug mold and plugs were incubated at 4 °C until solidified. For human peripheral blood mononuclear cells (PBMCs), peripheral blood of a healthy donor was isolated by density gradient centrifugation using Ficoll Paque Plus (GE Healthcare) according to manufacturer’s instructions. Plugs were prepared according to Plug Lysis protocol (BioNano Genomics). Briefly, 1×10^6^ cells were washed twice with PBS and mixed with 2% low melting agarose as described above. All plugs were incubated twice (2 hours incubation followed by an overnight incubation) at 50 °C with freshly prepared 167 μl Proteinase K (Qiagen) in 2.5 ml lysis buffer (BioNano Genomics) with occasional shaking. Next, plugs were incubated with 50 μl RNase (Qiagen) in 2.5 ml TE (pH 8) for 1 hour at 37 °C with occasional shaking. Plugs were washed three times by adding 10 ml wash buffer (10 mM Tris, pH 8, 50 mM EDTA), manually shaking for 10 seconds and discarding the wash buffer before adding the next wash. Plugs were then washed four times by adding 10 ml wash buffer and shaking for 15 min on a horizontal platform mixer at 180 rpm at room temperature. Following washes, plugs were stored at 4 °C in wash buffer or used for labeling. In order to extract high molecular weight DNA, plugs washed three times in TE pH 8 with shaking on a horizontal platform as explained, were melted for 2 min at 70 °C, followed by 5 min incubation at 43 °C. Next, 0.4 units of Gelase (Thermo Fisher) were added and the mixture was incubated for 45 min. Viscous DNA was gently pipetted and incubated at room temperature overnight in order to achieve homogeneity. DNA concentration was determined using Qubit BR dsDNS assay.

### Genetic barcoding and non-methylated CpG labeling

Genetic barcodes were produced by nick translation, a modified version of Irys prep NLRS protocol (Bionano Genomics). Briefly, 300 ng DNA were nicked with 10 units of Nt.BspQI (NEB) at specific sequence motifs (GCTCTTCN^) in the presence of 1 μl 10X buffer 3.1 (NEB) for 2 h at 50 °C and ultra-pure water to a volume of 10 μl. For DNA labeling, 15 units of Taq DNA polymerase were incubated with 200 nM of dATP dGTP dCTP (Sigma) and atto-532-dUTP (Jena Bioscience) in the presence of 1.5 μl of 10X thermopol buffer and ultra-pure water to a volume of 15 μl, the reaction was incubated at 72 °C for 1 h. Nicks were then ligated for 30 min at 37 °C using 4 units of Taq DNA ligase (NEB) in the presence of 0.5 μl thermopol buffer, 20 μM dNTPs (Sigma) 1 mM NAD+ (NEB) and water to a volume of 20 μl. For non-methylated CpG labeling (mTAG labeling) 300 ng of nick-labeled DNA were incubated overnight at 37 °C with 4.3 μg of the CpG-specific cytosine-C5 MTase M.SssI double mutant Q142A/N370A (eM.SssI)^26^ in the presence of 80 μM cofactor analogue AdoYnAzide^50^ and 3 μl 10X M.SssI reaction buffer (Thermo fisher/10 mM Tris-HCL, 50 mM NaCl, 1 mM DTT, pH 7.9), in a total reaction volume of 30 μl. The reaction was supplemented with 2 μM 5′-methylthioadenosine/*S*-adenosylhomocysteine nucleosidase (MTAN) and after 2 h of incubation, a spike-in of 1.1 μg of the eM.SssI and 2 μM MTAN^51,52^ was added to the reaction to further improve labeling efficiency. Next, 60 μg of Proteinase K (PK) were added and incubated for 2 h at 45 °C. Following PK digestion, 700 μM of dibenzocyclooctyne-cy5 (DBCO-cy5) were added and incubated overnight at 37 °C. The Dual-labeled DNA was then embedded in low melting agarose gel plugs and washed in 10 ml wash buffer (20 mM Tris-HCl, 50 mM EDTA, pH 8) for 15 min 5 times and twice with TE pH 8. Plugs were melted at 70 °C for 2 min and incubated at 43 °C for 5 mins. 2 μl of beta-agarase (Thermo Fisher Scientific) were added for agarose digestion and the reaction was incubated at 43 °C for 45 min. Dual-labeled DNA was stained with DNA stain (BioNano Genomics) according to the Irys Prep NLRS protocol with the addition of 25 mM Tris pH 8 and 30 mM NaCl. DNA concentration was measured by Qubit HS dsDNA assay.

To test non-methylation labelling efficiency, DNA was labelled with eM.SssI as described above, with or without MTAN. All DNA samples were purified from excess fluorophores, then applied on a custom epoxy-covered multi-well slide and imaged using commercial slide scanner, as described in the work of Margalit *et al*^53^.

### Glass slide preparation and quality-control imaging

22 × 22 mm^2^ glass cover-slips were cleaned for at least 7 hours to overnight by incubation in a freshly made 2:1 (v/v) mixture of 70% nitric acid and 37% hydrochloric acid. After extensive washing with ultrapure water (18 MΩ) and then with ethanol, cover slips were dried under a stream of nitrogen. Dry cover-slips were immersed in a premixed solution containing 750 μl N-trimethoxysilylpropyl-N,N,N-trimethylammonium chloride and 200 μl of vinyltrimethoxysilane in 300 ml ultrapure water and incubated overnight at 65 °C. After incubation, cover-slips were thoroughly washed with ultrapure water and ethanol and stored at 4 °C in ethanol. The silane solution was freshly made and thoroughly mixed before the cover-slips were introduced into the mixture. Stored cover-slips were normally used within 2 weeks.

For quality-control, samples were imaged to evaluate DNA length and degree of labeling. Samples were diluted 1:100–1:300 in TE with DTT buffer (10 mM Tris pH 8, 1mM EDTA, 200 mM DTT, Sigma) and were stained with 130 nM YOYO-1 (Invitrogen) DNA intercalating dye. DNA molecules were stretched by placing 8 μl of the solution at the interface of an activated coverslip placed on a standard microscope slide. The extended DNA molecules were imaged with a fluorescence microscope (TILL Photonics GmbH) using an Olympus UPlanApo 100X 1.3 NA oil immersion objective. Each image was composed of three colors, the YOYO-1, atto-532 and the Cy5 fluorophores, and was therefore imaged with the appropriate filters (485/20, 537/26 and 650/13 bandpass excitation filters, 525/30, 578/16 and 684/24 bandpass emission filters, for YOYO-1 and Cy5, respectively). Images were acquired by a DU888 EMCCD (Andor technologies) with an EM gain setting of 300 and exposure times of 100 for YOYO-1 and 2000 ms for atto-532 and Cy5.

### Global methylation quantification

Differences in global fluorescence intensity following labeling with and without MTAN, and between genomic samples were analyzed by an in house software^34,54^ (https://github.com/ebensteinLab/Tiff16_Analyzer). The code measures the total length of each molecule as well as the number and intensity of CpG labels detected along the DNA molecules. Quantification of non-methylated CpG was done by dividing the total intensity of the CpG label by the total length of the DNA (calculated as intensity of methylation signal / total DNA length in kbp). Similar analysis was performed on images of DNA stretched in nanochannels from CLL patient and healthy donor. In total we sampled 5.585 Gbp from CLL patient and 5.865 Gbp from the healthy donor and a total of 9840 field of view (FOV) from each donor.

### Optical mapping and analysis

Loading of DNA in nanochannels and imaging were performed using an Irys instrument (BioNano Genomics). Detection of imaged molecules and fluorescent labels along each molecule was performed by AutoDetect (version 2.1.4, BioNano Genomics). Alignment to the reference genome was performed using IrysView software (version 2.3, BioNano Genomics). *Arabidopsis* accession col-0 was run on a single chip on the Irys platform (BioNano Genomics), for up to 120 cycles to collect up to 89 Gb of quality filtered data. Molecule files (.bnx) were loaded into IrysView, quality filtered (>100 kbp length, >2.75 signal/noise ratio) and aligned to the reference TAIR10 (https://www.arabidopsis.org/download/index-auto.jsp?dir=%2Fdownload_files%2FGenes%2FTAIR10_genome_release).

Reference genome was converted into .cmap format and used to align the detected molecules using standardized parameters (-nosplit 2 -BestRef 1 -biaswt 0 -Mfast 0 –FP 1.5 -FN 0.15 -sf 0.2 -sd 0.0 -A 5 -outlier 1e-3 -outlierMax 40 -endoutlier 1e-4 -S -1000 -sr 0.03 -se 0.2 -MaxSF 0.25 -MaxSE 0.5 -resbias 4 64 -maxmem 64 -M 3 3 –minlen 100 -T 1e-6 -maxthreads 32 -hashgen 5 3 2.4 1.5 0.05 5.0 1 1 3 -hash -hashdelta 10 - hashoffset 1 -hashmaxmem 64 -insertThreads 4 -maptype 0 -PVres 2 -PVendoutlier - AlignRes 2.0 -rres 0.9 -resEstimate -ScanScaling 2 -RepeatMask 5 0.01 -RepeatRec 0.7 0.6 1.4 -maxEnd 50 -stdout -stderr –usecolor 1). Alignments were analyzed for genome coverage, and gaps using the .xmap output file.

Following alignment, data was uploaded to IrysExtract^38^ to generate an average intensity profile of fluorescence intensity across the genome. We used all molecules aligned with a confidence equal or higher than 8 (P<=10^−8^) and that had at least 70% of each molecule aligned to the reference. All regions in the genome with less than 15x coverage (3% of the genome), were excluded from the data. The analysis pipeline results in a genome-wide intensity bedgraph file at 500 bp resolution. The file allows further analysis and visualization in a genome browser as seen in Figures 4 & 5.

### Comparison of optical methylation mapping to bisulfite sequencing

Genome-wide bisulfite sequencing data (GSE43857)^36^ was processed to generate a bedgraph track of non-methylated CpGs represented as a “methylation score”. To calculate the methylation score, the non-methylated CpGs were padded +/−500bp and summed over 1 kbp bins to generate the bedgraph file. The score in each bin was then divided by the maximum score, setting the data between 0-1. The genome was then divided to 10 kbp windows and the mean score from each method (methylation score for the bisulfite data and relative intensity for the optical mapping data) was calculated for each window, K-S test was performed to test normality and since the data do not normally distribute, Spearman correlation analysis was performed using SPSS (IBM Corp. Released 2016. IBM SPSS Statistics for Windows, Version 24.0), to determine the linear dependence between bisulfite sequencing and methylation mapping.

### Analysis of CpG methylation distribution across gene body and siRNA

*pol-iv*-dependent 24 nt short interfering RNAs (siRNA) was retrieved from the previously published data^41^. miRNA primary transcript loci were extracted from miRBase ath.gff3 file (http://www.mirbase.org/ftp.shtml)^55^. *Arabidopsis* TAIR10 TAS loci were obtained from the ta-siRNA database^56^. The bed files of the 24nt siRNA loci, miRNA precursor loci, and TAS loci were used to measure the non-methylated CpGs from optical mapping and genome-wide bisulfite sequencing as detailed below (supplementary file 5).

First, a file containing all CpGs was created from TAIR10 FASTA by knickers (version 1.5.5 Bionano Genomics). Then, a non-methylated CpG track from genome-wide bisulfite sequencing data of *A. thaliana* was created using BEDTools (version 2.25.0) subtract^57^. Each site in the non-methylated CpG track was expanded by 500 bp on both sides to adjust to the optical mapping resolution. Genome coverage of non-methylated CpG was generated by BEDTools genomecov to receive a non-methylated CpG bedgraph. Finally, this file was uploaded to deepTools^58^ in addition to the average label intensity from optical mapping and the clusters bed file. Using computeMatrix in reference point mode, we computed the mean value for each bin from all regions 10 kbp upstream and downstream from the clusters. All clusters were set to position 0, and median coverage for the neighboring regions in 1 kbp bins was plotted.

For gene body analysis, we compared the average label intensity of non-methylated CpGs from optical mapping and genome-wide bisulfite sequencing data with a bed file containing all *A. thaliana* genes (TAIR10) (supplementary file 6). Using computeMatrix in scale-region mode, all genes were scaled to the same length in 100 bp bins. We added 1500 bp upstream and downstream to the TSS and TES (respectively) and computed the mean value for each bin.

### Safety Statement

No unexpected or unusually high safety hazards were encountered.

## ASSOCIATED CONTENT

Supporting information

Supplementary file 1-additional results as described in the main text (PDF)

Supplementary file 2-optical mapping intensity healthy blood sample.

Supplementary file 3-optical mapping intensity CLL sample.

Supplementary file 4-comparison between non- methylation CpG sites in optical mapping vs. bisulfite sequencing.

Supplementary file 5-24nt siRNA, miRNA precursor loci and TAS loci in *A. thaliana*.

Supplementary file 6-genomic locations of *A. thaliana* genes.

## AUTHOR INFORMATION

Y.E. and E.W. conceived the study. T.G performed optical mapping experiments and data analysis. Y.M, S.A and D.T performed experiments and data analysis. Z.S.Z provided MTAN enzyme. M.J and E.W provided M.Ssss I enzyme and cofactor. J.A.L and C.C. performed methylation analysis in Arabidopsis. N.A helped with human samples and clinical aspects. T.S, Y.E. and Y.M wrote the manuscript. All authors read and edited the manuscript.

## Corresponding author

*

## ACKNOWLEDGMENTS

Y.E. acknowledges European Research Council consolidator grant (817811). Y.E. and E.W. acknowledge funding from a joint Israeli German R&D Nanotechnology grant (Israel Innovation Authority, German Federal Ministry of Education and Research 13GW0282B and NATI 61976). Work in the Law laboratory was supported by an NIH NIGMS grant (GM112966) and a Salk Innovation Grant as well as the Rita Allen Foundation Scholars Program and the Hearst Foundation. C.C. was supported by a Postdoctoral Fellowship from the Glenn Center for Research on Aging at the Salk Institute and from a Salk Women and Science award. Z.S.Z was supported by NIH NIGMS (1R01GM101396)

We thank Kerstin Glensk for the preparation of eM.SssI. We thank Joseph R. Ecker and Florian Jupe for providing initial DNA samples.

