## Supplemental file 1- Supporting Figures 1-4 for "Chemoenzymatic labeling of DNA methylation patterns for single-molecule epigenetic mapping"

[1] School of Chemistry, Center for Nanoscience and Nanotechnology, Center for Light-Matter Interaction, Raymond and Beverly Sackler Faculty of Exact Sciences, Tel Aviv University, Tel Aviv, Israel.

[4] Department of Obstetrics and Gynecology, Meir Hospital, Kfar Saba, Israel & Sackler Faculty of Medicine, Tel Aviv University, Tel Aviv, Israel.

**Figure S1.** Quantification of non-methylated cytosine on labeled DNA molecules.

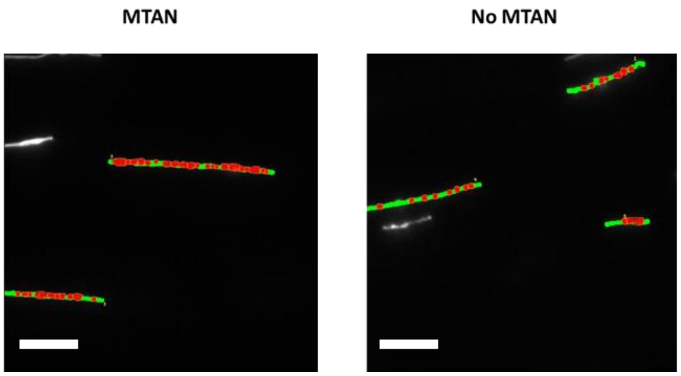


Representative image of unmethylated λ DNA stretched on activated glass slides and analyzed by an in-house developed software for measuring intensity profiles of colocalized non-methylated cytosine labels and stained DNA molecules. In green, YOYO-1 labeled molecules that were detected by the software. In gray, DNA molecules labeled with YOYO-1 that were rejected from the analysis due to non-linear stretching or crossing with another molecule. Labels at non-methylation CpG sites that were detected by the software are shown in red. Scale bars correspond to 20 kbp.

**Figure S2.** Labeling intensity profile measurement of eM.SssI, with or without MTAN.

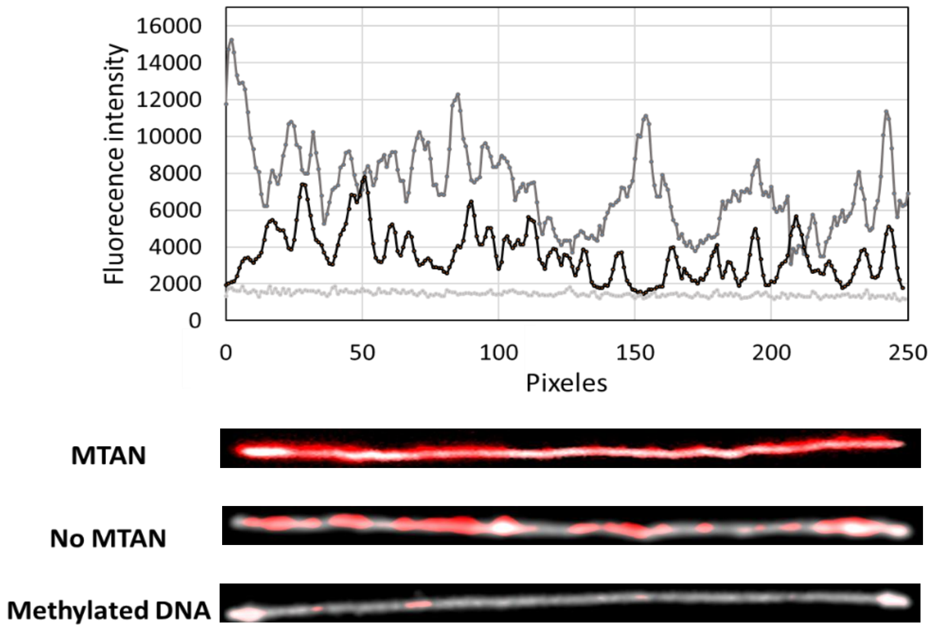


Top panel: Fluorescence intensity profile across unmethylated λ DNA, labeled with eM.SssI at CpG sites, with (dark gray) or without MTAN (black). Methylated λ was used as control (light gray). Bottom panel: Representative images of λ DNA molecules labeled with eM.SssI in the presence of MTAN on unmethylated DNA, in the absence of MTAN on unmethylated DNA and on methylated DNA. Here the DNA backbone is shown in gray and the labels marking non-methylated CpG sites are shown in red. Red images intensities were scaled for visualization.

**Figure S3.** Representative *in silico* representation of molecules aligned to chromosome 3.

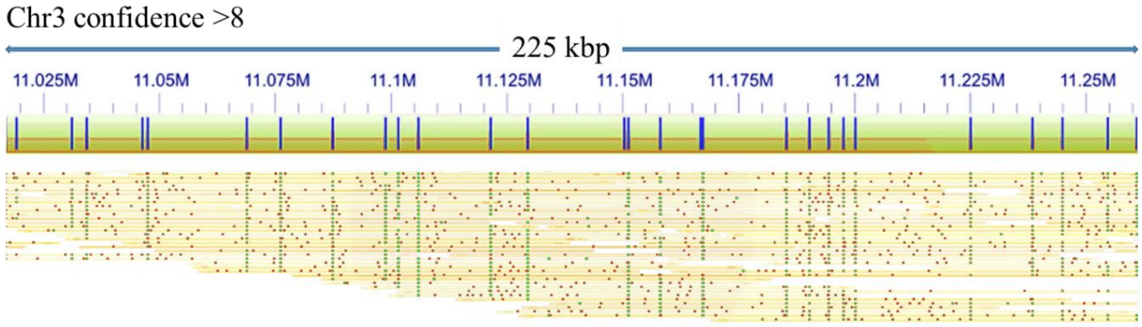


In green, genetic labels using Nt. BspQI, and in red, labels for non-methylated CpG sites marked by eM.SssI.

**Figure S4.** Non-methylated CpG sites over miRNA precursor and TAS loci.

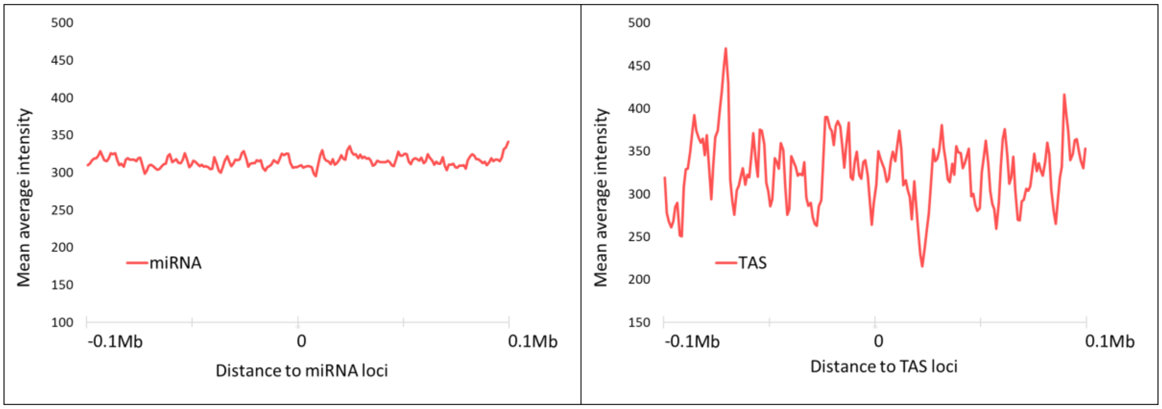


Optical mapping average intensity of non-methylated CpG sites across **A**. miRNA precursor loci **B**. TAS loci.
